# Visualisation and analysis of RNA-Seq assembly graphs

**DOI:** 10.1101/409573

**Authors:** Fahmi W. Nazarie, Barbara Shih, Tim Angus, Mark W. Barnett, Sz-Hau Chen, Kim M. Summers, Karsten Klein, Geoffrey J. Faulkner, Harpreet K. Saini, Mick Watson, Stijn van Dongen, Anton J. Enright, Tom C. Freeman

## Abstract

RNA-sequencing (RNA-Seq) is a powerful transcriptome profiling technology enabling transcript discovery and quantification. RNA-Seq data are large, and most commonly used as a source of genelevel quantification measurements, whilst the underlying assemblies of reads, if inspected, are usually viewed as sequence reads mapped on to a reference genome. Whilst sufficient for many needs, when the underlying transcript assemblies are complex, this visualisation approach can be limiting; errors in assembly can be difficult to spot and interpretation of splicing events is challenging.

Here we report on the development of a graph-based visualisation method as a complementary approach to understanding transcript diversity and read assembly from short-read RNA-Seq data. Following the mapping of reads to the reference genome, read-to-read comparison is performed on all reads mapping to a given gene, producing a matrix of weighted similarity scores between reads. This is used to produce an *RNA assembly graph* where nodes represent reads derived from a cDNA and edges similarity scores between reads, above a defined threshold. Visualisation of resulting graphs is performed using Graphia Professional. This tool can render the often large and complex graph topologies that result from DNA/RNA sequence assembly in 3D space and supports info rmatio no verlay on to nodes, e.g. transcript models. We have also implemented an analysis pipeline for the creation of RNA assembly graphs with both a command-line and web-based interface that allows users to create and visualise these data. Here we demonstrate the utility of this approach on RNA-Seq data, including the unusual structure of these graphs and how they can be used to identify issues in assembly, repetitive sequences within transcripts and splice variants. We believe this approach has the potential to significantly improve our understanding of transcript complexity.

## Introduction

The advent of next generation sequencing platforms enables new approaches to solving a variety of problems in medicine, agriculture, evolution and the environment. RNA-sequencing (RNA-Seq) based transcriptome analyses are now used routinely as an alternative to microarrays for measuring transcript abundance, as well as offering the potential for gene and non-coding transcript discovery, splice variant and genome variance analyses[1, 2]. Data are typically summarised by counting the number of sequencing reads that map to genomic features of interest, e.g. genes. These measures are used as the basis for determining the level of expression in a given sample and differential expression between samples. Currently, a large number of pipelines for the analysis of RNA-Seq data have been developed to go from the output of a sequencing machine, to sequence assembly, and on to the quantification of gene expression[3-5]. However, many aspects of the analysis of these data remain computationally expensive and limiting, and tools are still under active development[6-9].

One area in particular that remains challenging is the visualisation and interpretation of transcript isoforms from short-read data. There are a currently a number of software tools available for the visualisation of RNA-Seq assemblies, such as Manananggal[10], Integrative Genomics Viewer (IGV)[11],[12], Tablet[13], BamView[14], EagleView[15], Artemis[16], Vials[17], SpliceViewer[18], and JunctionSeq[19] (reviewed in [20]). Perhaps the most widely used sequence analysis and visualisation tool is the Integrative Genomics Viewer (IGV)[11, 12]. A distinguishing feature of IGV compared to other analysis/visualisation tools e.g. Tablet[13], BamView[14], EagleView[15] and Artemis[16], is the inclusion of the Sashimi transcript visualisation plot[21]. Sashimi produces plots for RNA-Seq derived transcript isoforms providing a quantitative summary of genomic and splice junction mapping reads together with gene model annotations and read alignments. Alignments in exons are represented as read densities and paired reads connect exons, where the connection is weighted relative to the number of reads aligning to a splice junction. Numerous other tools focus more on the quantitative and comparative visualisation of differentially expressed spliced exons and isoforms in different samples (reviewed by [20, 22]). In general, all existing visualisation approaches for splice variant analysis commonly involve the ‘stacking’ of reads onto a genomic reference. While this is sufficient for many needs, when the underlying transcript assemblies are complex and/or no reference genome is available, this approach can be limiting.

In DNA sequence analysis, *de novo* assembly methods based on the de Bruijn graph have proved to be highly effective in assembling a genome from millions of sequencing reads[23]. However, network visualisation of DNA sequence data has been little explored. Novák et al. (2010) first introduced the idea of using graph-based methods to visualise DNA assemblies, in this case repetitive sequences in plant genomes (pea and soybean) [24]. In this work, similarity scores between reads were pre-computed by a series of computationally intensive pair-wise sequence alignments, and represented the first step in building a network from such data[24]. After generating a matrix of similarity scores from an all-versus-all read comparison, read similarities exceeding a specified threshold were used to define network edges. In the visualisation of the assemblies, nodes represented individual reads of DNA sequence and edges denoted sequence similarity i.e. homology score between reads above a defined threshold. It was argued that the complex to po lo gy and diversity of the graphs produced could be used to better analyse the variability and evolutionary divergence of repeat families, as well a s to discover and characterise novel elements. Graph visualisations however were generated as PDF files, limiting the opportunity for data exploration. Nielson *et al.* (2009)[25] developed an interactive network display called ABySS-Explorer that allows a user to display a sequence assembly and associated meta-data. ABySS was developed to assemble sequencing data derived from individual human genomes. Here, the de Bruijn graph data structure of k-mer neighbourhoods was used to reduce memory usage and computation time. Contig assemblies are represented as an oscillating line, the number of oscillations corresponding to a fixed number of nucleotides. However, the graphs were not designed to support the visualisation of transcript splice forms.

Here we further explore the use of network-based visualisations of sequencing data, applying the fundamental principles first described by Novák *et al.* (2010)[24] to RNA-Seq data. Our aim has been to develop an approach that supports the visualisation of DNA assemblies, and in particular provides a means to better understand transcript structure and splice-variation. In this paper, we develop a novel method for the visualisation of RNA-Seq data using the graph analysis tool, Graphia Professional, formerly known as BioLayout *Express*^3D^ [26, 27]. In so doing, we provide a platform that supports the improved interpretation of complex transcript isoforms. We believe this approach will be useful in the exploration and discovery of new biological insights from sequence data.

## Methods

### RNA-Seq data

Four samples of cell-cycle syncronised of RNA-Seq data were generated from serum-starved human fibroblasts (NHDF) [28]. Briefly, cells were starved for 48 h and then harvested at 0 h and following serum refeeding at 12, 18 and 24 h, as the cells underwent synchronised cell division. RNA-Seq analysis was performed on an Illumina HiSeq2500 (Illumina, San Diego, California, USA) with 100 b ppaired-end sequencing according to the manufacturer’s recommendations and performed by Edinburgh Genomics (Edinburgh, UK) using the TruSeq™ RNA Sample Prep Kit (Illumina). Poly-(A) RNA was isolated and fragmented to produce of an average 180 bp fragments. Fragmented RNA was reverse transcribed and a single stranded DNA template was used to generate double strand cDNA which was blunt ended using T4 DNA polymerase prior to the addition of an adenosine base to assist ligation of the sequencing adapters. Flow cell preparation was carried out according to Illumina protocols; the libraries were denatured and diluted to a concentration of 15 pM for loading into the flow cells. RNA-Seq data were processed using the *Kraken* pipeline, a set of tools for quality control and analysis of high-throughput sequence data[29]. Expression levels were reported as Fragments Per Kilobase of transcript per Million (FPKM).

To analyse a broader range of samples, RNA-Seq data from a human tissue atlas[30] representing 27 different tissues were downloaded from ArrayExpress database (E-MTAB-1733). Primary visualisation of the data was performed using IGV to visualise the reads mapped on to the reference genome in certain loci or genes across samples. Long-read sequencing data of human heart, liver and lung samples released by Pacific Biosciences (PacBio)[31] were also utilised for making comparison to transcript assembly generated from the short read data.

### Preparation of files for transcript visualization

The pipeline described below is based around a set of linked bash and Python scripts that perform the following tasks. Initial QC and read mapping to the reference genome (GRCh38) were performed using *BowTie* v1.1.0 [32]. Sequence mapping data (BAM) were converted to a text file suitable for graph visualisation using Graphia Professional (Kajeka Ltd, Edinburgh, UK). Firstly, BAM files were sorted according to mapped chromosomal location using *sort* from *SAMtools*[33]. The R package *GenomicRanges*[34] was used to create annotation information, from a GTF file containing node annotation. This GTF file holds annotation information about gene structure (Ensembl version GRCh38). The output from this step was a tab-delimited file containing read mappings on Ensembl transcript and exon features. This information can be overlaid on to graphs using the *class sets* function of Graphia, such that upon selection of an Ensembl transcript ID, nodes representing reads that map to this transcript model will be coloured according to the exon number.

The next step was to define the similarity between reads mapping to a gene of interest from the BAM and GTF files. A FASTA file containing all sequences mapping to a particular gene was extracted and the supporting information used for the visualisation of transcript isoforms in the context of the resultant graph. For read-to-read comparison MegaBLAST[35] was used to generate a similarity matrix with edge weights derived from the alignment bit score. Parameterisation of this step i.e. defining the threshold for % sequence similarity (*p*) and length (*l*) over which two sequences must be similar in order for an edge to be drawn between them is of particular importance. Ideally, a graph should contain the maximum number of reads (nodes), connected by a minimum number of edges and where possible give rise to a single graph component i.e. a single group of connected nodes that together represent the mRNA species of interest. For high coverage transcripts, more stringent parameters may be desirable.

### Graph layout

Graphia Professional originally used a modified version of the Fruchterman-Reingold (F-R) algorithm [36] for graph layout. Whilst the existing implementation of the F-R algorithm was capable of producing layouts for many large biological networks, the unusual topology of DNA/RNA sequence graphs neccessitated a new and highly optimised graph layout approach. In order to test available alternative layout algorithms, an idealised overlap network was generated by assuming 100 ordered reads (nodes) where each read overlaps the previous read by 95% of its length, i.e. edge weight between adjacent nodes is 0.95. In this paradigm the first read would overlap successive reads by 5% less each time, sharing only 5% similarity to read 20 and no similarity to subsequent reads. A second file of this type was prepared so as to represent two splice variants, where one variant was identical to the first network but a second variant included a 50 node addition where similarities started to branch off after the first 50 nodes, rejoining the graph at position 51. Following examination of these synthetic ‘transcript’ graphs a series of tests were performed using data mapping to collagen type 1 V alpha 1 (*COL5A1*), a long (8.5 kb) and highly expressed gene in human fibroblasts encoded by a single transcript species. This was used to evaluate the performance of the layout algorithm on a large RNA assembly graph. Following experimentation, the Fast Multipole Multilevel Method (FMMM)[37] was shown to be well suited to the layout of these types of graphs. The FMMM algorithm was reimplemented in Java from the Open Graph Drawing Framework (OGDF)[38] and incorporated into the Graphia code base, adding uniquely the ability to perform graph layout in 3D space.

### Collapsing of redundant reads

In the case of highly expressed genes there can be a significant degree of redundancy in read coverage i.e. reads of exactly the same sequence may be present in the data many times. This makes the read-to-read comparison step unnecessarily time-consuming and the resultant graph sometimes difficult or impossible to visualise due to its size, whilst redundant reads add nothing to the interpretation of transcript structure. Using *Tally* from the *Kraken* package, multiple identical reads were mapped to a single identifier that incorporates the number of occurrences of that specific sequence. In the visualisation, a single node was used to represent multiple identical reads and the diameter of a node was proportional to the original number of occurrences of the reads it represents.

### Analysis of the graph structure

Initially, we chose to examine a set of 550 genes whose expression was up-regulated as fibroblasts entered into S-M phases of the cell cycle (18-24 h after being refed serum). A graph derived from the 24 h data was plotted for each gene using MegaBLAST parameters *p*=98, *l*=31. Where the topology of a given gene graph was relatively simple, an explanation of its structure required only the overlay of individual transcript exon information in order to identify splice variant(s) represented. In other cases more detailed analyses were required. Other graphs were generated from the human tissue atlas data available at ArrayExpress (E-MTAB-1733)[30]. In the tissue samples, reads may originate from multiple cell types expressing different isoforms of the same gene. The 100 bp paired-end reads for each tissue were individually mapped to the human genome (Ensembl GRCh38.82) with STAR v2.3.0[39]. The output from the mapping process (BAM files) was used to generate RNA assembly graphs using our pipeline. Publicly available data for *TPM1* were used to compare the network-based RNA-Seq approach with Pacific Biosciences (PacBio) long-read results obtained through their website[31]. The *TPM1* gene models from both data were compared for heart, brain and liver.

### Validation of splice variants using RT-PCR

To validate the existence of splice variants predicted by graph analyses, reverse transcription polymerase chain reaction (RT-PCR) of candidate splice variants was performed. Total RNA from human fibroblasts used for the RNA-Seq experiment was reverse transcribed in order to generate single stranded cDNA. Primers were designed using the Primer3 software[40] to amplify the region for validation of the splice variant. For *LRR1*, a pair of primers was designed to amplify three splice variants as suggested from the graph visualisation while for *PCM1* two pairs of primers were designed across two different splice variant locations. For *LRR1*: Forward primer 5’-TGTTGAGCCTCTGTCAGCAG-3’ and reverse 5’-GTGTGGGCAACAGAATGCAG-3’; for PCM1 (primer set 1) Forward primer 5’-TCTGCTAATGTTGAGCGCCT-3’ and reverse 5’-TGCAGAGCTAGAAGTGCAGC-3’ and *PCM1*: (primer set 2: Forward 5’-ACGGAAGAAGACGCCAGTTT-3’ and reverse 5’-AGCTGCAGCTCATGGAAGAG-3’. PCR was carried out for 35 cycles (92°C, 30 sec; 60°C, 90 sec; 72°C, 60 sec). The amplicons were run on a 2% agarose gel in the presence of SYBR-Safe DNA gel stain (Thermo Fisher, Waltham, MA, USA) and gels visualised by UV illumination.

### Access to the pipeline and web-based NGS Graph generator

Documentation and full source code for the NGS Graph Generator package can be downloaded from: https://github.com/systems-immunology-roslin-institute/ngs-graph-generator. The user needs to supply BAM and GTF files to run the pipeline. In addition, we have developed a web interface that allows the pipeline to be run on a number of supplied datasets. This web-interface is called *NGS graph generator* and can be accessed at http://seq-graph.roslin.ed.ac.uk. Using this resource, a user can select a BAM file from RNA-Seq time-course samples of human fibroblasts or data from the human tissue atlas. Users can adjust parameters used by MegaBLAST to compute read similarity. There is an option to uniquify the graphs, i.e. remove redundant reads. Processing time required is dependent on the number of reads mapping to a gene of interest. The user must provide their email address and will be informed once the job finished. The resultant graph layout file will automatically open Graphia Professional (if installed). Protocols for graph generation and visualisation are provided in Supplementary File 1.

## Results

The principle of rendering sequence data as a network is illustrated in Figure 1A. We have developed a pipeline to create such graphs from RNA-Seq data, where the outputs can be visualised as a graph within Graphia Professional (Figure 1B). For these studies, we generated RNA-Seq data from four samples of human fibroblasts, taken at different time points during synchronised cell division. Paired-end cDNA libraries for each RNA sample were prepared and sequenced (see Methods). Other RNA-Seq data used for these studies originated from a publicly available human tissue atlas experiment [30] (E-MTAB-1733) which was also processed through the pipeline.

**Figure 1:**
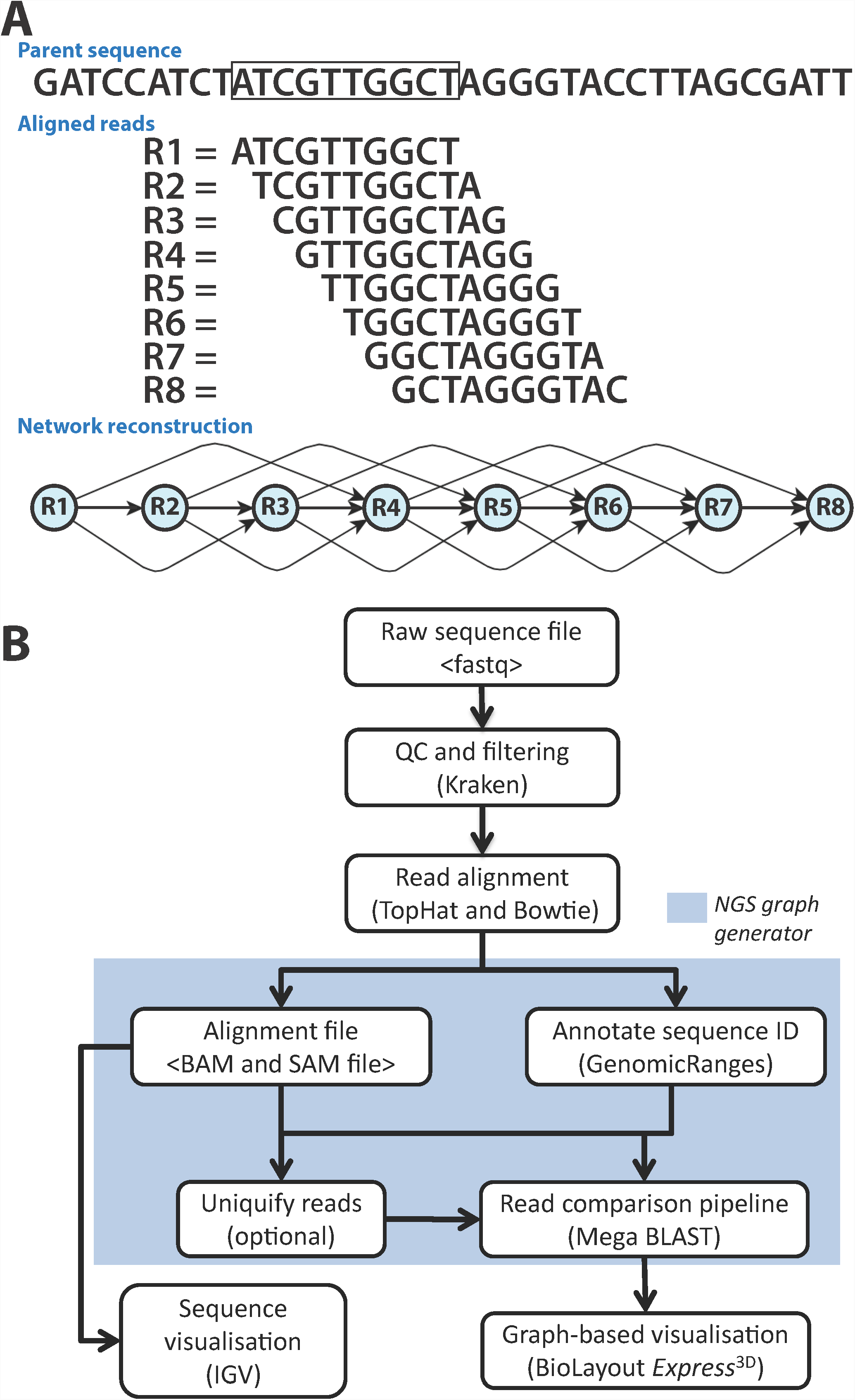
Development of the analysis pipeline for visualisation of RNA-Seq data. **(A)** Network paradigm for the analysis of short-read RNA-Seq data. A region of DNA is shown with 10 bp ‘reads’ aligned to it below. In this view, eight different reads position with a 1 bp consecutive offset to the parent sequence. If reads were compared and a threshold of >70% similarity were used to construct a graph, the network would have structure shown. **(B)** Pipeline for network-based visualisation of RNA-Seq data. Analysis pipeline for building graphs from RNA-Seq data passes from raw sequencing FASTQ file through a series of analysis/annotation steps up to the production of a file for graph visualisation within Graphia Professional.

### Optimisation of graph visualization

An optimal layout is crucial to the interpretation of graph structure. Initial studies of RNA-Seq graph visualisation demonstrated that the Fruchterman-Reingold (F-R) algorithm originally used by Graphia Professional [36] for graph layout performed poorly on these types of graphs (Figure 2). Following examination of the available algorithms for graph layout we incorporated the fast multipole multilevel method (FMMM)[37] algorithm into the tool, enabling FMMM graph layout in a 3D environment. The implementation of FMMM provides an interface where alternative force models can be selected (F-R or Eades) and includes various settings that offset layout quality versus the speed of graph layout. Graph visualisation of a theoretical matrix of 100 reads where consecutive reads overlap by 95% demonstrates the difference between the F-R and FMMM algorithms (Figures 2Ai-v). Graphs take on a corkscrew-like appearance at a local level when the layout algorithm is set to a ‘high quality’ mode, i.e. it employs more iterations of the FMMM algorithm (Figures 2Aiii-v). Indeed, the more edges present (defined by the stringency of similarity cut off) the tighter a graph is coiled. This unique feature of overlap graphs can also be observed in graphs generated from RNASeq data, particularly when the depth of sequencing is high. We then explored the layout of RNA-Seq data (from NHDF 24 h post-serum refeeding; see Methods) for *COL5A1*; a 66 exon, 8,471 bp long transcript that is highly expressed by fibroblasts (40,170 reads mapped to this gene in this sample). Figure 2Bi shows the layout of the *COL5A1* RNA assembly graph using the F-R layout algorithm, contrasted to its layout using the FMMM algorithm (Figure 2Bii). In general, the higher the FMMM quality setting, the more linear a graph becomes, but at the cost of computational runtime.

**Figure 2:**
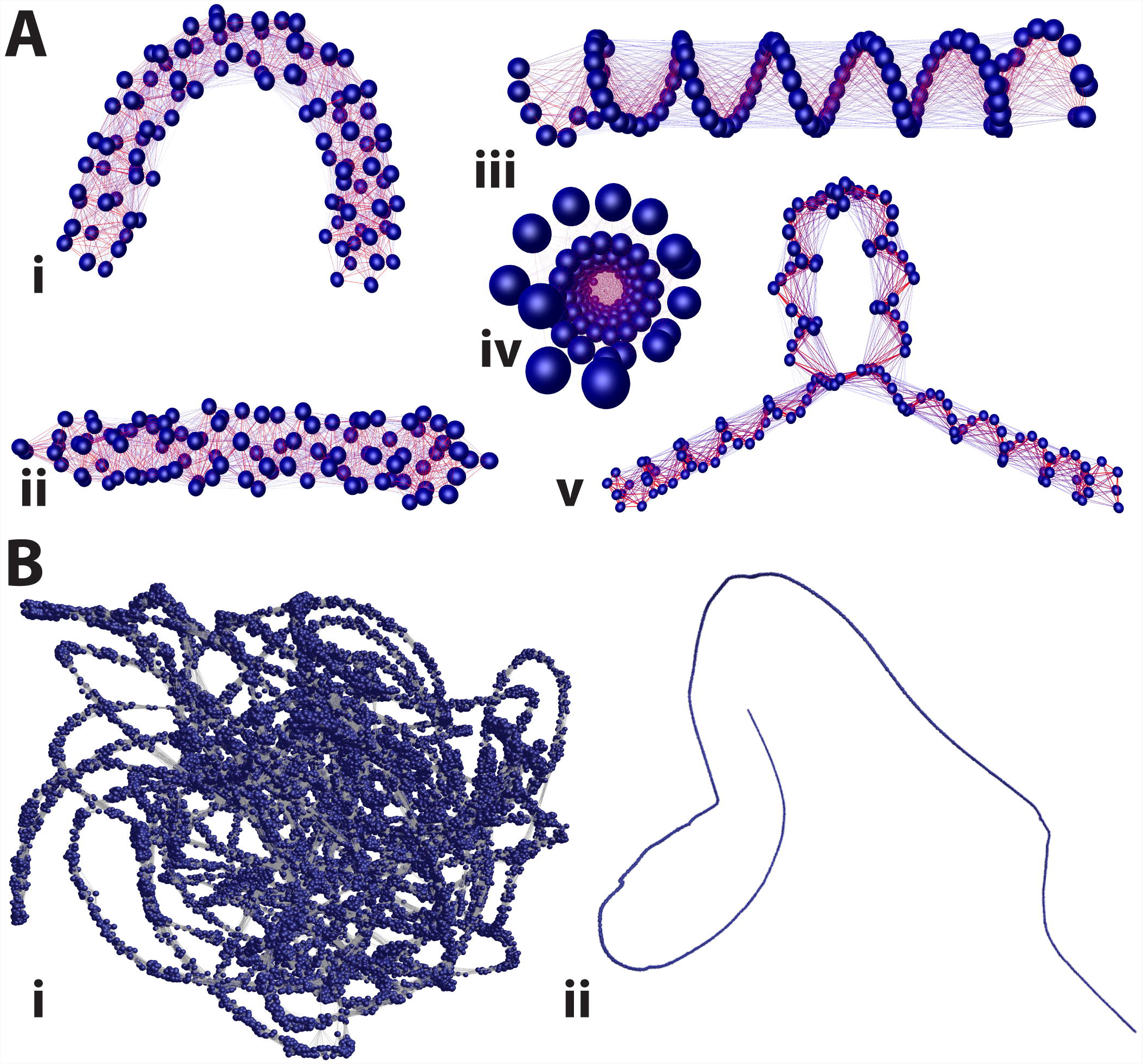
Optimisation of graph layout. **(A)** Layout of a perfect overlap matrix consisting of 100 ‘reads’ where consecutive reads overlap by 95% and a ≥5% similarity has been used as minimum threshold for defining edges. In example **Ai** a modified Fruchterman-Reingold layout was used to layout the graph, in **Aii** the FMMM algorithm using the Eades force model and a ‘Low Quality, High Speed’ setting and **Aiii** is the same as **Aii** but the ‘High Quality, Low Speed setting’ was used. **Aiv** is an end on view of **Aiii** illustrating the corkscrew-like structure of the graph. **Av** is a visualisation of an alternatively spliced transcript generated from simulated data. **(B)** Graph visualisation of *COL5A1*, which is a highly expressed gene in human fibroblasts. **Bi** shows the graph layout using the Fruchterman-Reingold algorithm as originally implemented within Graphia Professional and **Bii** the layout of *COL5A1* using FMMM algorithm (Eades force model and the High Quality, Low Speed setting).

### Graph complexity reduction

Another factor in defining graph structure is the criteria used to define the similarity score between reads. The two MegaBLAST parameters that define sequence similarity scores based on the length and percent identity (*l* and *p*) are adjustable. If the thresholds for these settings are too high, graphs may fragment and fine structure will be lost; if too low the underlying structure may be obscured and it will greatly increase the memory footprint of a graph and layout time due to an excess number of edges. After empirical exploration of the two variables for thresholding the read-to-read similarity score using a range of RNA assembly graphs, we chose a percentage similarity *p=98* and percentage length coverage *l=31,* as reasonable generalised values for these variables.

In some instances where the level of expression is especially high, graph visualisation is not possible due to the number of nodes and edges needed to represent the data. For instance, in the 24 h serum-refed fibroblast samples the highly expressed genes *TUBA1C* and *GAPDH* had 38,294 and 59,998 reads mapping to them, respectively. Node reduction is a process whereby identical reads are collapsed down to and represented by a single node, the size of the node being proportional to the number of nodes it represents. In the case of *TUBA1C*, this reduced the number of nodes from 38,294 to 6,511 (Figure 3A), whilst the number of edges was reduced from 90,340,179 to 1,779,069. In the case of *GAPDH*, the reduction in nodes was from 59,998 to 9,264, whilst the number of edges was reduced from 208,221,932 to 3,562,688 (Figure 3B). The reduced graph for *GAPDH* is also shown generated at two different MegaBLAST threshold settings. On the left the graph was generated at the default BLAST setting of *p*=98, *l*=31, the second used a more stringent setting of *p*=98, *l*=95. Such is the depth of sequencing of this gene that even using a BLAST setting of 98% similarity over 95 b pof length the graph still forms a single component where the number of edges is reduced by approximately 90% but the number of nodes by only about 1%. At this higher stringency BLAST setting the graph uncoils exposing small nodes representing unique reads due to sequencing errors (as shown in inset of Figure 3B).

**Figure 3:**
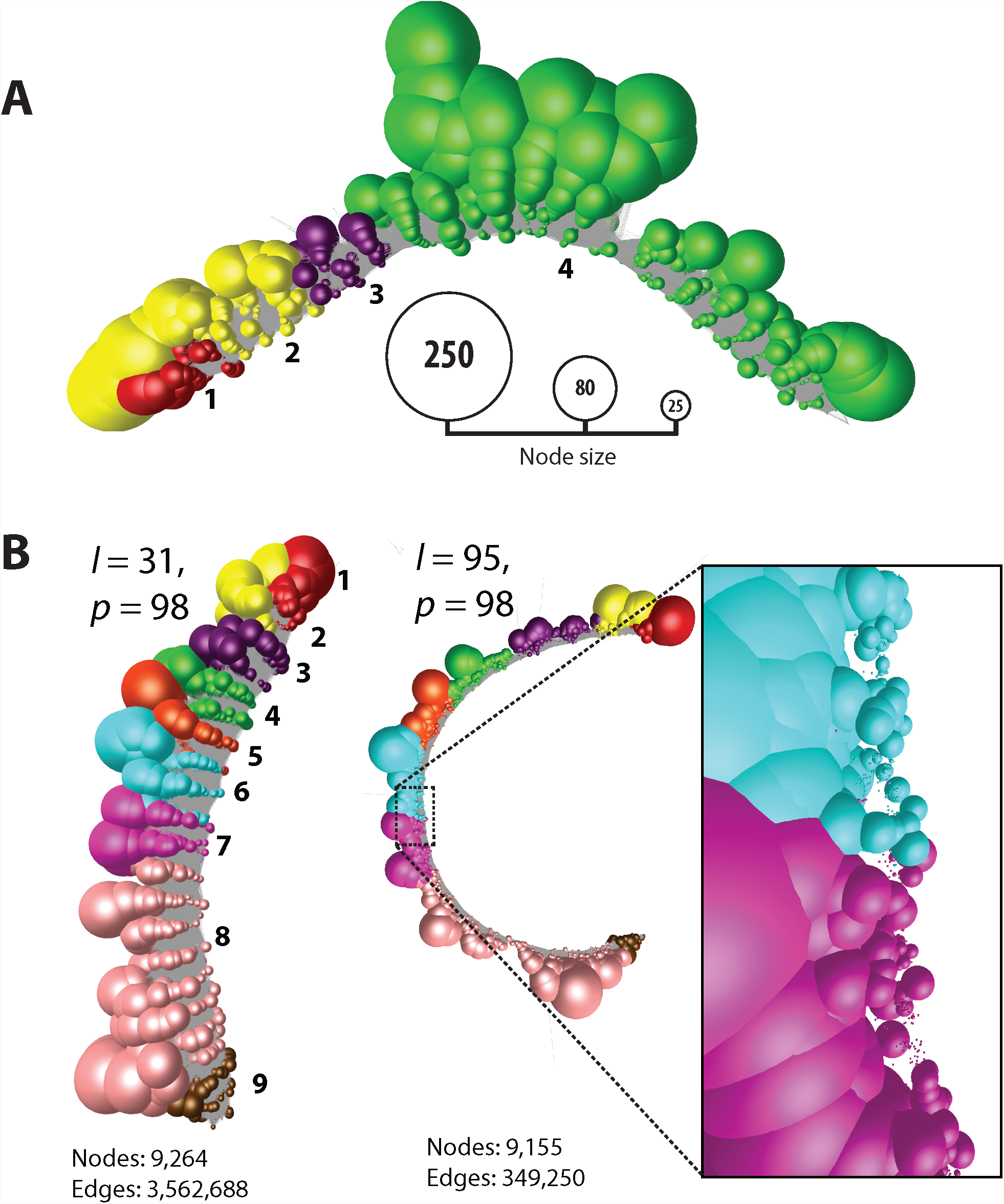
Read uniquification of highly expressed genes TUBA1C **and** GAPDH **in human fibroblasts.** **(A)** Graph representation of *TUBA1C* after read uniquification. The number of reads mapping to *TUBA1C* was 38,294 in the fibroblast sample. After read uniquification this was reduced to 6,511 unique reads. When graphs are collapsed down to unique sequences node size is proportional to the number of individual reads represented, and nodes have been coloured according to the exon on to which they map. The *TUBA1C* transcript model matching the graph here is ENST00000301072, a 3,001 bp transcript encoding a 449 amino acid protein. **(B)** Graph for *GAPDH* after read uniquification as described above but shown at two different thresholds of BLAST scores. On the left using our default threshold of 98% similarity (*p*) over 31 bp (*l*), on the right 98% similarity over 95 bp. Increasing the edge threshold (*l*) dramatically reduces the number of edges, whist the number of nodes is barely affected. It also opens up the structure and when edges are removed the large number of small nodes representing unique reads due to sequencing errors can be observed (inset). The *GAPDH* transcript model matching the graph here is ENST00000396856, a 1,266 bp transcript encoding a 260 amino acid protein.

### Network visualisation of transcripts

Having optimised general settings for network construction, we individually examined 550 RNA assembly graphs from the set of genes up-regulated as NHDF cells undergo mitosis. In the majority of cases, the graphs for these genes were linear, i.e. possessed no higher order structure; two example transcripts, *KRT19* and *CCNB1*, from two time points (0 h and 24 h post-serum) are shown in Figure 4. Arguably little has been learnt by visualising these data in this manner. However, even with these networks transcript variance could be observed. The effect of a change in expression level could be inferred from a reduction/increase in the number of nodes. In the network representing *CCNB1* there was clear evidence that two transcript isoforms were expressed by the fibroblasts, one of which was truncated at the 3’ end. This was manifest by the fact that there were fewer reads present at the 3’ end of the network, the falloff in read density occurring at the point where a known variant occurs (ENST00000505500), and the coiled structure broke down beyond that point (Figure 4iii). This decrease in reads at the 3’ end of *CCNB1* is also visible in the standard IGV Sashimi plot (Figure 4iii).

**Figure 4:**
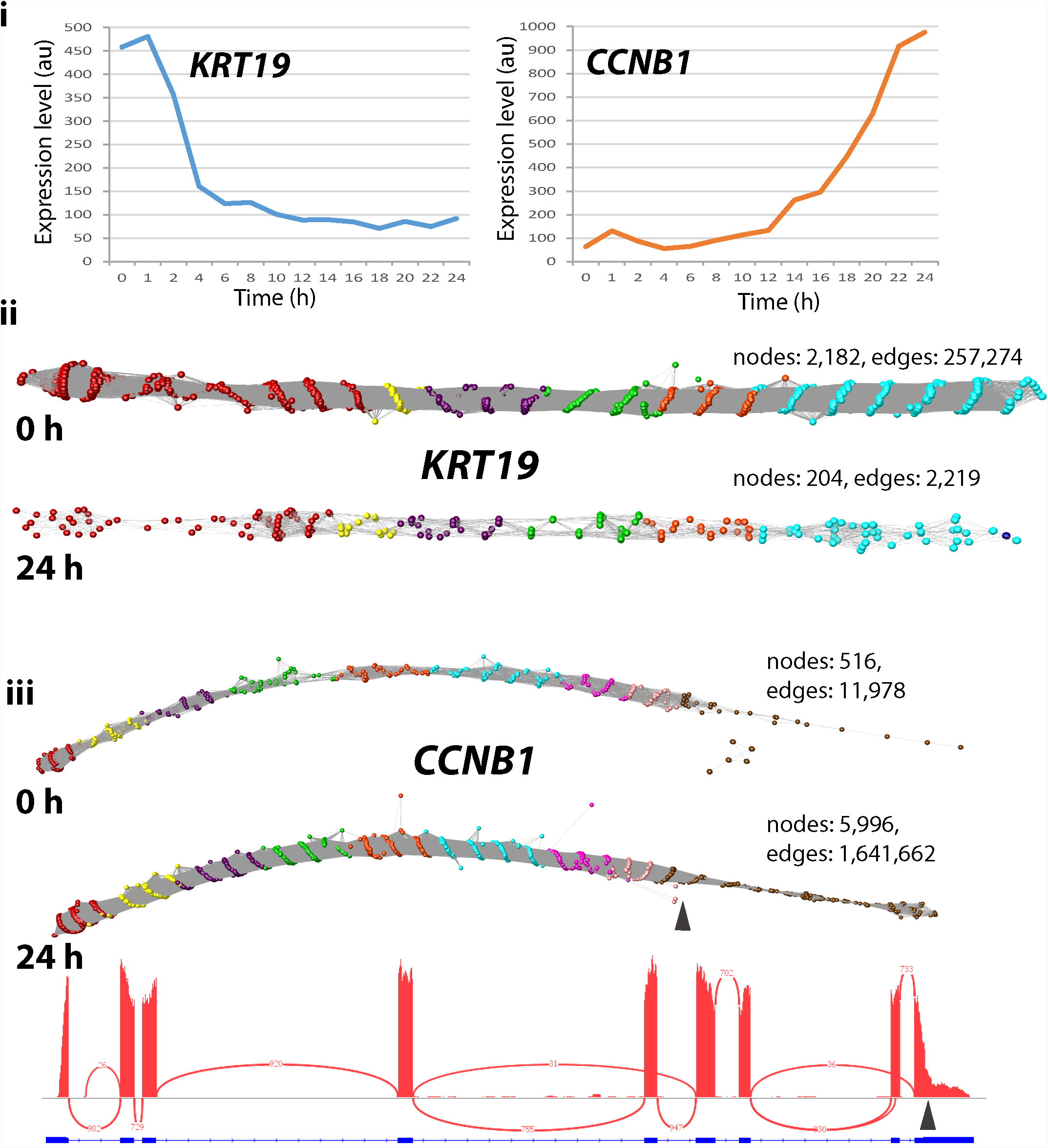
Typical graphs of RNA-Seq data derived from linear transcripts. Examination of a wide range of RNA-Seq assembly graphs derived from human fibroblast expressed genes demonstrated that the majority of graphs are linear structures. Shown here are two such graphs for *KRT19* and *CCNB1* (node overlay colours representing exons were derived from ENST00000361566 and ENST00000256442, respectively). Top: expression profile of the two genes as measured by microarray analysis of the time-course of transcriptional events following serum-refeeding. Expression of *KRT19* is rapidly down-regulated whilst *CCNB1* is up-regulated as the cells enter mitosis. This differential expression is evident from the graph visualisation with the number of nodes decreasing or increasing by approximately 10 fold in the 0 h derived vs. the 24 h derived RNA-Seq data. It is interesting to note that in the *CCNB1* graphs there is an abrupt decrease in the density of nodes within exon 9 at both time points (marked by arrow). This corresponds to where the IGV view also shows a decrease in the density of reads and corresponds to a *CCNB1* transcript (ENST00000505500) that exhibits a truncated exon 9 at this position.

### Splice variant network structure

Of the 550 differentially expressed transcripts examined in the NHDF data, approximately 5% of RNA gene assembly graphs exhibited complex topologies. We investigated the underlying reasons for these unusual structures. LRR1 (leucine-rich repeat protein 1) is known to regulate the cell cycle in *C. elegans* and actin-based motility in human cells[41]. The *LRR1* graph comprised of a single loop-like structure and corresponded to the two known transcript isoforms for this gene. One (ENST00000318317) has only three exons, where exons 3 and 4 are skipped, while a second protein coding transcript (ENST00000298288) contains four exons, numbered 1, 2, 4 and 5 (Figure 5A). There was also evidence for the presence of a nonsense mediated decay product (ENST00000554869) as a small number of reads partially mapped to a small exon (numbered exon 3) specific to this transcript. *PCM1* (pericentriolar material 1) is a 6,075 nt gene containing 36 exons that encodes a protein that recruits PLK1 to the pericentriolar matrix to promote primary cilia disassembly before mitotic entry[42]. There are seven known protein coding variants of PCM1 and network analysis provided evidence for two splicing events when expressed in fibroblasts (Figure 5B). One loop was indicative of the splicing out of exon 7 and the other of exon 24, indicating the presence of transcripts ENST00000517730 and ENST00000522275 respectively, in addition to the main isoform of this gene (ENST00000325083). RT-PCR confirmed the splicing events for *LRR1* and *PCM1* genes predicted by the network-based analysis (Figure 5A iv and B ii). The visualisation of splice variants was also supported in the Sashimi plots for these genes, but even in these relatively simple examples of splice variation the plots can be challenging to interpret, especially in the case of *LRR1* (Figure 5Aiii).

**Figure 5:**
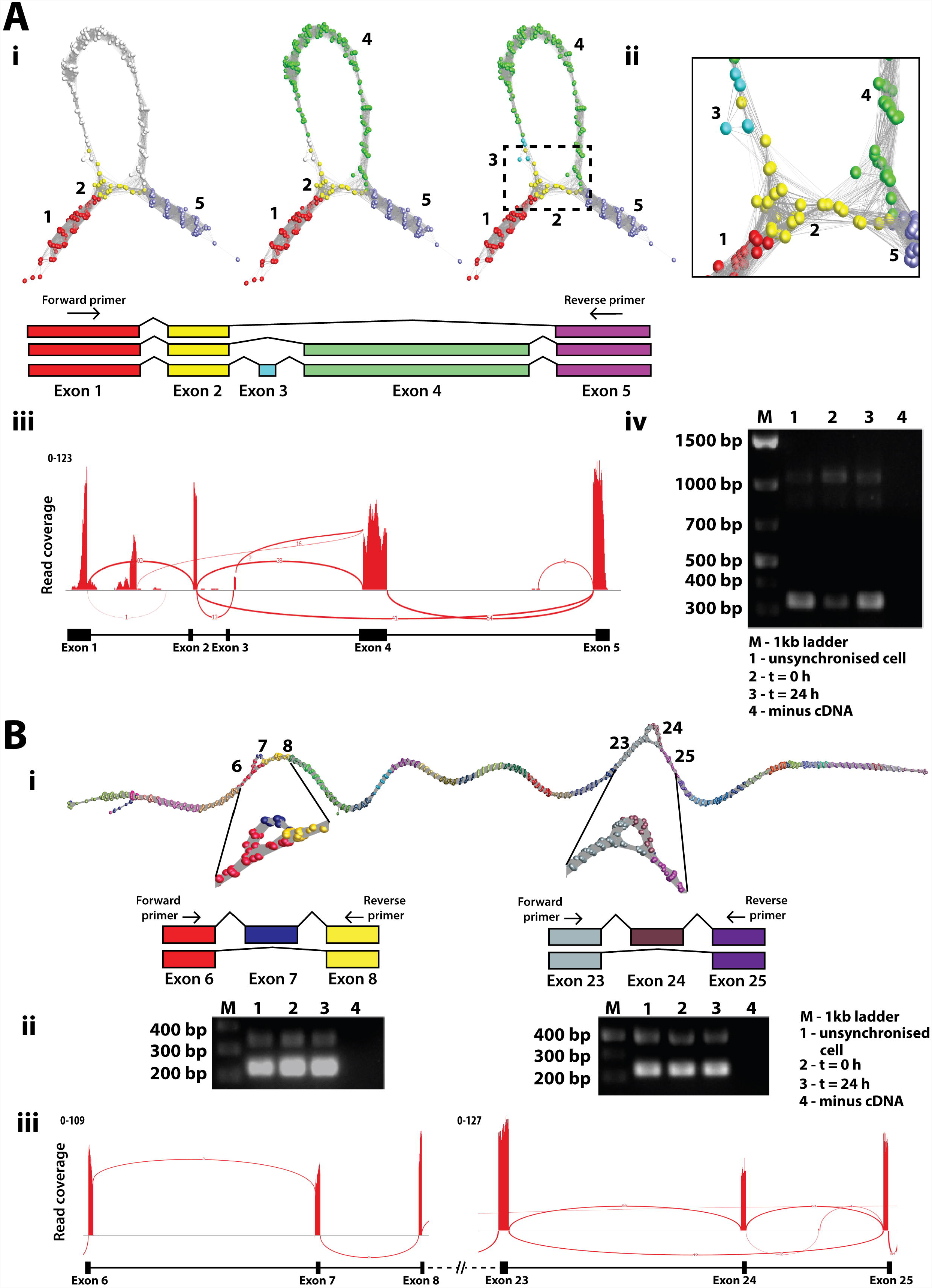
Splice variant visualisation and confirmation. **(A)** Splice variants of *LRR1*. **Ai** Loop in the *LRR1* graph suggested that two or possibly three transcript isoforms are expressed in fibroblasts. These are shown overlaid on the graph together with a schematic representation of each isoform. **Aii** Close-up view of exon skipping event in *LRR1* shows the connection of exon 2 (yellow nodes) to exon 5 (light purple nodes) skipping exons 3 and 4 (blue/green nodes). **Aiii** Sashimi plot generated in IGV showing RNA-Seq reads mapping to the *LRR1* locus. Splice junctions are displayed as arcs connecting exons. The number of reads observed for each junction is indicated within segments, and y-axis ranges for the number of reads per exon base are shown. **Aiv** Result for RT-PCR of *LRR1* mRNA using the human fibroblast RNA from the proliferation time course. Three bands showing on the gel represent alternatively spliced products due to exon 3/4 skipping (310 bp), exon 3 skipping (1022 bp) or the full transcript (1130 bp). The position of the PCR primers is shown in **Ai. (B)** Splice variants of *PCM1*. **Bi** Two different splicing events for *PCM1* were evident from the graph visualisation of this gene. **Bii** Result for RT-PCR of *PCM1* for two different locations of splice variant using the human fibroblast RNA from the proliferation time-course. Skipping of exon 7 resulted in a PCR fragment size of 223 bp, compared with 339 bp when included, and skipping of exon 25 resulted in a PCR fragment of 331 bp compared with 496 bp when included. **(iii)** Representative Sashimi plot generated in IGV.

### Observations of issues with assembly and internal repeats

CENPO (Centromere O protein) is a component of the Interphase Centromere Complex (ICEN) and localises at the centromere throughout the cell cycle[43], where it is required for bipolar spindle assembly, chromosome segregation and checkpoint signalling during mitosis. The RNA assembly graph of *CENPO* showed a complex topology within its final 3’ exon (Figure 6A). In principle, network elements representing single exons should form linear graphs with bifurcations from linearity only occurring at exon junctions. In order to explain the observed anomalies in the graph for this gene, we investigated the genomic origin of reads mapping to loop junctions using BLAST. It transpired that many mapped to exon-exon junctions of adenylate cyclase 3 (*ADCY3*), a gene located on the opposite strand of chromosome 2. A number of its 5’ exons overlap with the final 3’ exon of *CENPO*. The RNA-Seq libraries were non-directional in nature and due to an ambiguity in read mapping, reads in the assembly of *CENPO* were actually derived from *ADCY3*. Reads from exon boundaries of *ADCY3* gave rise to the observed alternative splicing-like structures within the final portion of the *CENPO* graph. These anomalies are difficult to observe using conventional visualisation tools such as IGV and even with the Sashimi plot it is not easy to distinguish which of the two overlapping genes reads are derived from.

*MKI67* encodes antigen Ki-67, a well-established cell proliferation marker, that helps support the architecture of the mitotic chromosome[44]. The graph of *MKI67* contains twofeatures; a loo prepresenting a known splice variant, and a knotted structure associated with exon 14. In the case of the former, exon 7 is spliced out in transcript ENST00000368653 as compared to ENST00000368654, both isoforms being expressed within fibroblasts. In the second case there are 13 repeats of a K167/chmadrin domain within exon 14, the internal homology leading to the formation of the observed structural complexity (Figure 6B).

**Figure 6:**
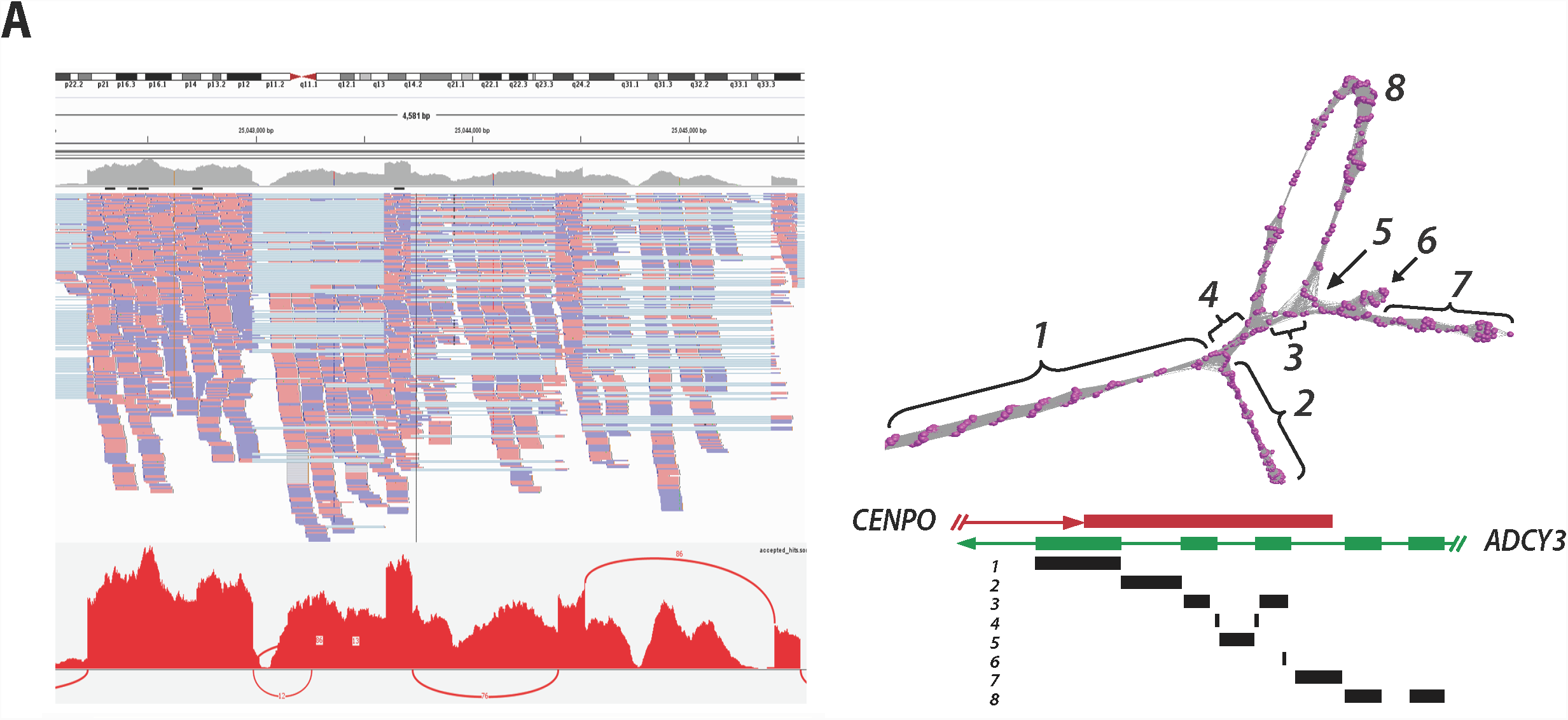
Complex gene graph structure. **(A)** Mis-assembly of reads from overlapping genes. **Ai** IGV visualisation of *CENPO* exon 8 together with corresponding Sashimi plot. *ADCY3* overlaps with *CENPO* on the opposite strand of DNA. **Aii** In this case reads derived from *ADCY3* mRNA are being wrongly mapped to *CENPO* resulting in the complex graph structure observed in exon 8. The schematic diagram of overlapping genes *CENPO* and *ACDY3* is shown above and regions of the graph mapped back to it. The loops in exon 8 of *CENPO* are formed by junction reads derived from *ADCY3* encoded on the opposite strand. **(B)** Repeat sequences cause perturbation in graph structure. Network-based visualisation of *MKI67*. In this graph, there are two structures: **(1)** an alternatively spliced exon and **(2)** internal duplication. Skipping of exon 6 giving rise to ENST00000368653 can be observed as the loop structure, whilst the knotted structure is formed due the presence of 16 K167/Chmadrin repeat domains within exon 12.

### Human tissue graph analysis of *TPM1* gene

In order to explore transcript variation within and between tissues we examined expression patterns across a human tissue atlas. We focus here on *TPM1*, a gene that encodes the muscle/cytoskeletal protein alpha tropomyosin. This gene was selected because it is widely expressed across tissues but at very variable levels, and was identified by the *rMATS* package[45] as having high statistical probability of containing multiple splice variants between the tissues examined, i.e. heart, liver, brain. It has 15 exons and Ensembl reports there to be 19 protein coding transcript isoforms, with a further 14 non-coding isoforms, i.e. with a retained intron or the products of nonsense mediated decay (see http://www.ensembl.org). We analysed graphs of *TPM1* generated from three different human tissues (heart - 10,347 reads; liver - 431 reads; brain - 1,006 reads) using public human RNASeq data (see Methods). In each case the RNA-assembly graph demonstrated varying degrees of complex topology (Figure 7A).

**Figure 7:**
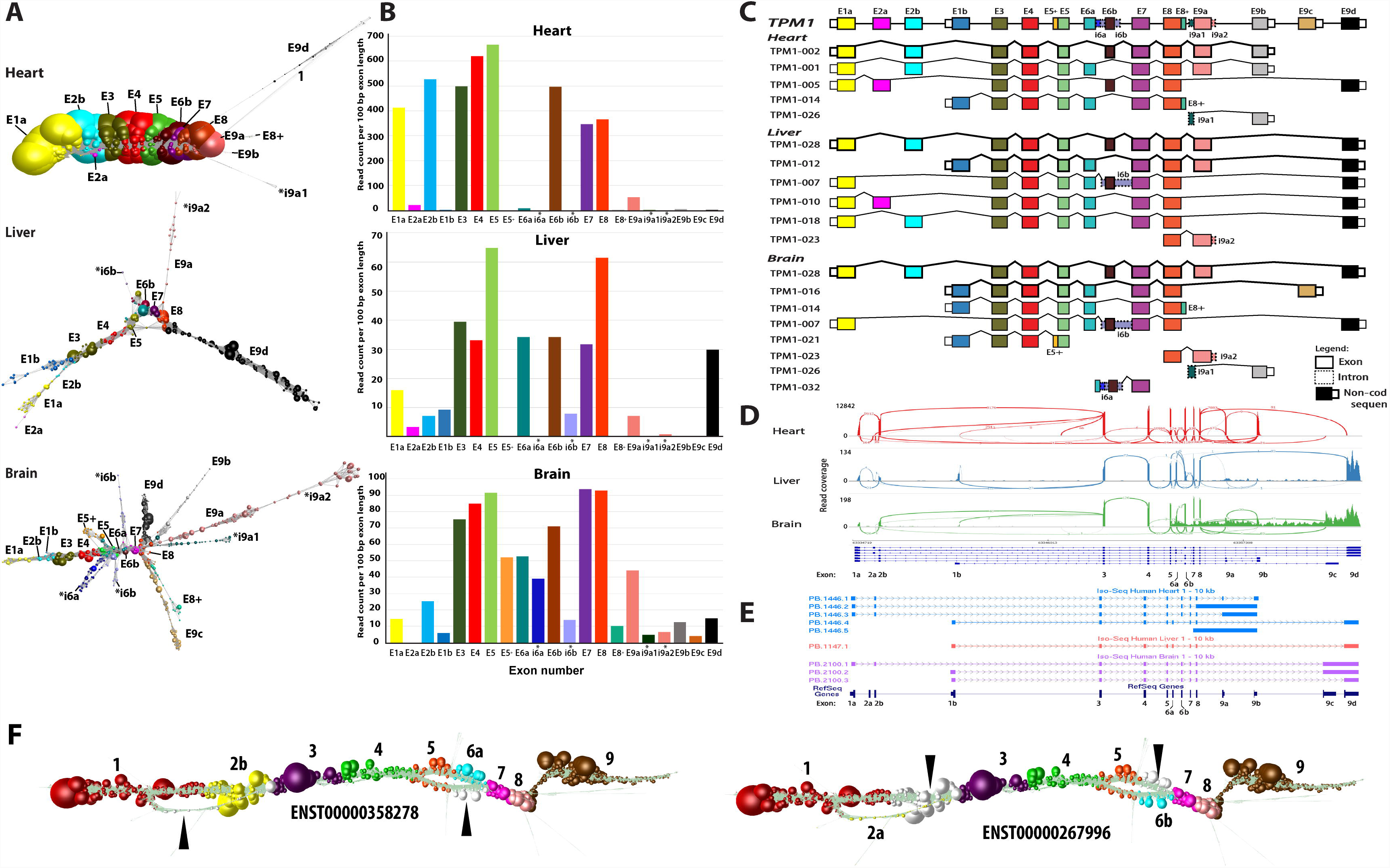
Graph visualisation of TPM1 transcript isoforms expressed in different tissues. **(A)** Graphbased visualisation of *TPM1* mRNA from heart, liver and brain. In these graphs, node size reflects the relative read depth as shown in Figure 3. In the case of the heart, the *TPM1* expression graph shows one major isoform while an estimated four other minor isoforms are observed that give rise to network ‘branches’. In the *TPM1* graph generated from liver, all isoforms would appear to be expressed at the similar levels. Alternative 5’ transcript initiation can be visualised as the first two branches of the graph. The mutually exclusive exons 6a and 6b can be seen from the graph structure, while the retention intron of 6b and 9a is observed as a few nodes branching out from the core graph structure. The RNA assembly graph generated from brain demonstrates that multiple isoforms are expressed in this tissue, a complex series of branched structures emanating out from the graph at different locations. The observed structure is supported by Ensembl gene models. **(B)** Histogram of the number of reads per 100 bp of exon per sample. **(C)** Schematic representation of *TPM1* transcript isoform diversity expressed in heart, liver and brain as interpreted from RNA assembly graphs. **(D)** Representative Sashimi plots generated in IGV showing RNA-Seq reads mapping to *TPM1* locus in the different tissues. Height of the bars represents read coverage. Splice junctions are displayed as arcs, with the number of reads observed for each junction being indicated by line thickness. **(E)** UCSC Genome Browser sequence visualisation of *TPM1* gene from a different study of the whole human transcriptome of brain, heart and liver generated on the PacBio long read platform. The number of isoforms detected by PacBio is different from the graph-based based visualisation. PacBio detected five isoforms in heart (blue), one in liver (orange) and three in the brain (purple). **(F)** *TPM1* graph as expressed in fibroblasts. Two bifurcation events are clearly visible in the graph that correspond to two isoforms of the transcript, ENST00000358278 and ENST00000267996. These exhibit two mutually exclusive exon splicing events, the former containing exons 2b and 6a and the latter containing exons 2a and 6b, the white nodes (arrows) corresponding to reads that do not map to the corresponding transcript.

Graph analysis of *TPM1* expression in the heart revealed that there was essentially only one major transcript isoform expressed in this organ (ENST00000403994) containing exons 1a, 2b, 3, 4, 5, 6b, 7, 8 and 9a/b, the deep coverage of which can be inferred from the size of nodes. However, there was evidence of the presence of other minor transcript isoforms, visualised as small branches emanating from the core network. Mapping the reads from these minor structures to *TPM1* exons suggested the expression of up to four other minor transcript isoforms in the heart, based on the presence of isoform-specific exons. In contrast the *TPM1* assembly graph derived from liver suggests at least six isoforms expressed in this organ. The two major isoforms (ENST00000559556, ENST00000404484)possess alternative 5’ ends as can be observed from the bifurcation point at the 5’ end of the graph (Figure 7A). An alternative splicing event at exon 6 can also be inferred from the graph visualisation and two 3’ termini are visible, both major isoforms ending in exon 9d and a minor form ending in exon 9a. There was also evidence of retained intronic sequence, shown by reads mapping to intron 6b. The most complex *TPM1* graph was derived from the brain. We observed what we believe to be eight isoforms expressed in brain tissue. There appear to be three major isoforms (ENST00000559556, ENST00000317516, ENST00000560975) and others showing evidence of retained introns. These retained intronic sequences can be inferred from the branching structures, e.g. introns 5, 6 and 8. The major splice isoforms are at the 3’ end at exon 9c and 9d. Another isoform can be inferred from the nodes branching at exon 8. Two isoforms in which intron 9a1 or exon 9b were retained were expressed in this sample. We compared the network-based analyses of *TPM1* as summarised in Figure 7C with the corresponding Sashimi plots (Figure 7D) and data from PacBio long-read sequencing of cDNA tissues from these tissues[31] shown as UCSC tracks (Figure 7E). In the case of the Sashimi plot we would argue that it is very difficult to work out the structure of the expressed transcripts from this visualisation and whilst PacBio the long read result agrees with our analysis, it misses much of the complexity *TPM1* isoform expression. Finally, we looked at the expression of this gene in the fibroblast data. In this instance two isoforms of the transcript were observed, ENST00000358278 and ENST00000267996 which exhibited two mutually exclusive exon splicing events, the former containing exons 2b and 6a and the latter containing exons 2a and 6b (Figure 7F). The *TPM1* example explored here demonstrates something of the potential complexity of alternative splicing and where RNA assembly graphs may help with data interpretation.

## Discussion

RNA-Seq offers a platform to explore transcript diversity within and between cells and tissues. Sequencing platforms that produce short-read data (50-250 bp) currently dominate the field. Many tools and analysis pipelines already exist to process these data from the DNA sequencer, through mapping to a genome or *de novo* assembly, and summarise these data down to read counts per gene/transcript. These data are then ready for differential gene expression or cluster-based analyses. It is also routine practice to port data into tools such as IGV, where they can be visualised in the context of the reference genome (where available). Reads are shown stacked on to the region from which they have been determined to originate. In instances where a single RNA species is transcribed from a specific locus, existing visualisations are sufficient for most needs. However, where multiple transcripts are produced from a given locus, deconvolution of that assembly into the component transcripts can be challenging. Tools such as the Sashimi plot use information derived from exon boundaries and paired end reads to display connections between exons, the thickness of the joining line indicating the number of reads derived from a given exon-exon boundary. When transcript diversity is relatively simple these views provide a sufficient representation of events, but when transcript variability is complex they can be difficult to interpret. Other tools for splice variant analysis focus on identifying statistically splice variants from data and whilst some possess sometime sophisticated visualisations, they generally rely of the same read stacking approach to display transcript isoforms.

This work describes a complementary approach to the analysis and interpretation of RNA-Seq data, based on construction and visualisation of RNA assembly graphs. In this method RNA-Seq reads mapping to a specific locus are directly compared with each other by calculating an all-vs-all similarity matrix. In the context of a graph visualisation of these data, nodes represent individual reads or collections of identical reads, whilst edges represent similarity scores between them, above a given threshold. Information about a read can be used to annotate nodes. In this manner different transcript models can be overlaid easily on to the graph assemblies, reads derived from a given exon sharing the same colour. This provides a way to quickly visualise how well a given transcript model matches the structure of the assembly.

A fundamental challenge in implementing this approach is the ability to layout and display graph assemblies of data, rapidly and such that the underlying structure of the networks can be interpreted. Novak *et al.*[24] visualised their graph assemblies of plant genomic repeat elements as image files, but these provide only an overview of a graph’s gross topology and no ability to interact or easily overlay additional information. Graphia Professional, formerly known as BioLayout *Express*^3D^, was originally developed for the visualisation and analysis of pathway models[46] and gene expression data as correlation networks[26, 27]. In this latter paradigm graphs can consist of tens of thousands of nodes (each representing a gene or transcript) connected by millions of edges (correlations), and for this reason the tool was designed to support the visualisation and exploration of very large graphs. Correlation networks often exhibit complex topologies with regions of high connectivity being due to groups of coexpressed genes. RNA assembly graphs however have a fundamentally different type of structure. Reads generally only share edges with others derived from up or downstream positions in the genome, thereby giving rise to a string-like topology. The FMMM algorithm uses a combination of an efficient multilevel graph layout scheme and a strategy for approximating the repulsive forces in the system, by rapidly evaluating potential fields[37]. When used at its highest quality setting to layout linear RNA assembly graphs, at a local level nodes form a corkscrew-like structure, whilst at a gross level they straighten out, allowing clear visualisation of the underlying structure. Lower quality but higher speed settings of the algorithm are often sufficient to appreciate higher level graph structure with greatly reduced layout times.

Here we explore the power of graph visualizations of RNA-Seq read assemblies to interpret transcript diversity. When a sole transcript is expressed in the data, a linear string-like graph is generated. If the coverage of the gene is low, the network may appear as a number of isolated components, whilst at higher levels of coverage a single graph component is formed. As coverage increases the graph starts showing a characteristic spiral appearance as a ‘perfect’ overlap graph is achieved. Up to a point, the depth of sequencing for a given transcript is immediately obvious from its graph visualisation. In two cases shown of *KRT19* and *CCNB1* (Figure 4) the decrease and increase in their expression is reflected in the density of nodes in the resulting graphs, as well as the alternative splicing at the 3’ end of the latter. At the point where every base position along a transcript has the start of one read mapping to it, in principle further reads add nothing to an assembly graph’s structure. Additional reads also add greatly to the computation time taken to calculate a read similarity matrix, and more nodes and edges have to be rendered. Collapsing redundant reads to a unique sequence/node therefore speeds up all aspects of visualisation. Currently the analysis pipeline only simplifies down to unique reads, so reads that contain a polymorphism or sequencing errors are represented as separate nodes. Abundant identical reads are represented by large nodes, the size of the node decreasing with read depth. The graph assemblies for *GAPDH* and *TUBA1C* (Figure 3) are used to illustrate this approach. These graphs represent single transcripts sequenced at high depth, the many small nodes representing a visualisation of sequencing ‘noise’. The number of edges is largely determined by the threshold set for read similarity. In order to minimise layout times and improve the fluidity of graph rendering, a similarity threshold should ideally be selected that allows the construction of a single component network with a maximum number of nodes and a minimum number of edges. This ‘ideal’ threshold value is dependent on the depth of sequencing, i.e. number of reads per unit length of DNA. From our experiments optimising MegaBLAST settings (Figure 3B) we defined a sequence similarity (*p*) < 0.02, and *l* value of 31% as a default thresholds – to potentially be made more or less stringent depending on read coverage.

Whenever an RNA assembly graph takes on higher order structure it is likely that sequences diverge, contain homologous domains, or there is an issue with sequence assembly. Where more than one transcript from the same gene is expressed in a sample, forks or loops in the graph are observed starting and finishing at exon boundaries. *LRR1* as expressed in human fibroblasts is a relatively simple example of where two alternatively spliced transcripts are expressed by the same cell population. In one version of the *LRR1* transcript expressed by fibroblasts, exon 4 is spliced out and a large loop is observed in the graph. The graph of *PCM1* possesses two loops corresponding to known splice variants at exons 7 and 24, both validated here. These splicing events are immediately obvious from the network visualisations, but perhaps less easy to appreciate from the corresponding Sashimi plot. In certain gene graphs we observed intra-exonic secondary structure. In the case of *MKI67* a series of 14 K167/Chmadrin domains present within exon 14 of the gene cause a knotted structure in that portion of the graph. An alternative splice variant missing exon 6 is also apparent in the graph visualisation of this gene. Within exon 8 of *CENPO* we observed complex network topology. In this case our analyses showed that it was due to reads produced from the transcription of *ADCY3* whose terminal 5’ exons overlap on the opposite strand. These mis-mapped exon boundary reads from the transcripts of *ADCY3* caused loops within graph representing exon of *CENPO* (Figure 6A). The inability to correctly map reads from over-lapping transcribed exons is one of the reasons the majority of RNA-Seq analyses are now generated from directional cDNA libraries.

We next wanted to test the potential of graph visualisation to analyse transcript complexity within and between tissues. In the case shown here, we examined *TPM1* (tropomyosin 1) transcript diversity in RNA-Seq data derived from three human tissues, heart, liver and brain. *TPM1* is highest expressed in the heart (and other muscles) where it functions as an actin-binding protein involved in the contractile system of muscles [47]. The dominant and possibly the sole functional transcript isoform was expressed in heart, although a relatively small number of reads mapped to exon 2a and to terminal intronic sequences, suggesting the presence of a low number of other transcript isoforms. Whether these represent transcriptional noise or transcription of these isoforms by cell types present in a low abundance in the heart is not clear. Expression levels of *TPM1* in the liver and brain are approximately 10 times lower than in the heart. In these tissues TPM1 is thought to play different roles in cytoskeletal organisation[47]. The two RNA assembly graphs generated for *TPM1* exhibited complex topologies. Through studying these graphs and mapping this information to the Ensembl transcript models for this gene, we estimated up to 6 transcript isoforms to be expressed in the liver, versus 10 in the brain. This is largely based on the presence in the data of reads mapping back to transcript-specific exons. Even with the availability of graph visualisations and other tools, interpreting these data was a challenge as transcript assemblies such as these are inherently complex. We found that the publicly available PacBio analyses of these three tissues was in partial agreement with our analyses. Although we detected all the isoforms identified through the PacBio data, many additional transcript isoforms were suggested by our analysis and not reported in the PacBio data. *TPM1* transcripts expressed by fibroblasts were different yet again, showing two mutually exclusive exon splicing events. It is interesting to note the different graph topologies of an exon skipping versus a mutually exclusive exon splicing event, the former giving rise to a loop such as in the case of *LRR1* or *PCM1*, the latter a bifurcation and rejoining of the graph as seen with *TPM1* expression in fibroblasts.

Here we present a new and complementary approach to aid the analysis of RNA-Seq data. We describe an analysis pipeline whereby reads mapping to given loci are compared for sequence homology and assemblies visualised as a graph. In this environment, information can be overlaid on the graph in terms of node colour and/or size (potentially shape) and the structure of the network can be explored in 3-dimensional space. Graph structure reveals the nature of the underlying sequence assembly and complexities therein. We demonstrate the ability to recognise splice variants in these graphs, as well as areas with internal homology and issues with read mapping using this approach. We also provide an example of just how complex these assemblies can be and the strengths and limitations of graphs and other approaches in these instances. The visual cues provided by this graph visualisation can be used to explore the reasons behind observed complexity and complement other solutions to the analysis of these data.

One of the limiting factors of this approach is the requirement to know in advance which portions of the data might be worth visualising as an RNA assembly graph. Hence, we recommend that other tools are used to detect splice variants such as DEXSeq[48], Cuffdiff[49], JunctionSeq[19] or rMATs[45] prior to network construction. Another limitation currently, is the inability to examine more than one RNA assembly graph/gene at a time. It can however be envisaged that assemblies could also be further collapsed such that many transcripts could be viewed as simplified graphs allowing large portions of sequencing data to be rendered. We are currently working on tools where even bigger graphs can be visualised and edge thresholds can be filtered dynamically, to enable more rapid and detailed analyses of RNA sequencing assemblies. In addition, we certainly see potential for this approach to be used to analyse transcript assemblies and DNA contigs where no reference genome is available, and options for visualisation are limited. In summary, we believe the data pipeline, the tools and basic approach presented here provide an effective analytical paradigm that is a novel contribution to the analysis of the huge amounts of information-rich but complex data produced by modern DNA sequencing platforms.

## Acknowledgements

This work was funded by the Biotechnology and Biological Sciences Research Council (BBSRC) grant no. BB/J019267/1 and T.C.F is funded by an Institute Strategic Grant from the Biotechnology and Biological Sciences Research Council (BBSRC) (BB/JO1446X/1). F.W.N. was funded by Majlis Amanah Rakyat (MARA), a Malaysian Government agency.

## Author contributions

T.C.F. conceived of the idea, supervised the project, and together with A.J.E. applied for the grant that funded the work. F.W.N. performed much of the explorative work described and helped in the development of the pipeline together with G.J.F., H.K., M.W., S.vD., A.J.E. and M.W.B. and S-Z.C. generated the NHDF data used in much of these analyses. T.A. and K.K. worked to the implement and modify the FMMM graph layout algorithm and F.W.N., B.S., K.M.S., S.vD., A.J.E. and T.C.F. wrote and edited the manuscript.

## Figures

**Supplementary file 1. Protocol for the generation and visualisation of RNA assembly graphs.** Description of pipeline and protocols for generating graphs using the NGS Graph Generator website, Unix, Amazon cloud and for analysing the results.

